# Interference of Neuronal TrkB Signaling by the Cannabis-Derived Flavonoids Cannflavins A and B

**DOI:** 10.1101/2022.02.03.478734

**Authors:** Jennifer Holborn, Alicyia Walczyk-Mooradally, Colby Perrin, Begüm Alural, Cara Aitchison, Adina Borenstein, Jibran Y. Khokar, Tariq A. Akhtar, Jasmin Lalonde

**Author notes:** Correspondence to: Jasmin Lalonde x. 54706. J.H. and A.W.M. contributed equally to this report.

## Abstract

Cannflavins A and B are flavonoids that accumulate in the *Cannabis sativa* plant. These specialized metabolites are uniquely prenylated and highly lipophilic, which, *a priori*, may permit their interaction with membrane-bound enzymes and receptors. Although previous studies found that cannflavins can produce anti-inflammatory responses by inhibiting the biosynthesis of pro-inflammatory mediators, the full extent of their cellular influence remains to be understood. Here, we studied these flavonoids in relation to the Tropomyosin receptor kinase B (TrkB), a receptor tyrosine kinase that is activated by the growth factor brain-derived neurotrophic factor (BDNF). Using mouse primary cortical neurons, we first collected evidence that cannflavins prevent the accumulation of Activity-regulated cytoskeleton-associated (Arc, also known as Arg3.1) protein upon TrkB stimulation by exogenous BDNF in these cells. Consistent with this effect, we also observed a reduced activation of TrkB and downstream signaling effectors that mediate *Arc* mRNA transcription when BDNF was co-applied with the cannflavins. Of note, we also performed a high-throughput screen that demonstrated a lack of agonist action of cannflavins towards 320 different G protein-coupled receptors, a result that specifically limit the possibility of a TrkB transinactivation scenario via G protein signaling to explain our results with dissociated neurons. Finally, we used Neuro2a cells overexpressing TrkB to show that cannflavins can block the growth of neurites and increased survival rate produced by the higher abundance of the receptor in this model. Taken together, our study offers a new path to understand the reported effects of cannflavins and other closely related compounds in different cellular contexts.

## 1. INTRODUCTION

Flavonoids are polyphenolic compounds found in various plant-derived foods and beverages. These phytochemicals represent a large family of molecules that can be classified into six main subclasses, based on their chemical structure: flavonols, flavanols (also known as flavan-3-ols or catechins), flavanones, flavones, anthocyanins, and isoflavones (Panche et al., 2016). Interestingly, evidence suggests that moderate habitual intake of flavonoids can lower the risk of cardiovascular disease, cancer, as well as all-cause mortality (Bondonno et al., 2019). Another purported benefit of these natural products is their positive influence on brain health and function, as several members of the flavonoid family have been found to promote neuroprotection, reduce neuroinflammation, and enhance cognition (Vauzour et al., 2008; Jaeger et al., 2018; Bakoyiannis et al., 2018). More precisely, flavonoids appear to modulate signaling pathways that are central to the control of neuronal survival and plasticity, such as the MAPK-CREB and PI3K-mTOR cascades (Vauzour et al., 2008; Jaeger et al., 2018). However, this particular line of inquiry has been little studied and therefore represents an opportunity to identify new bioactive compounds with therapeutic qualities among this family of phytochemicals.

Apart from the psychoactive molecule Δ^9^-tetrahydrocannabinol (THC) and other related cannabinoids with only mild or no psychotropic effect, like cannabidiol (CBD) and cannabigerol (CBG), the *Cannabis sativa (C. sativa)* plant also produces hundreds of specialized metabolites including at least twenty different flavonoid compounds (Flores-Sanchez et al., 2008). Among those, the flavones cannflavin A and cannflavin B (Figure 1A) are considered to accumulate uniquely in *C. sativa* cultivars. Seminal work by Barrett and colleagues performed more than 30 years ago helped identify these two flavonoids and characterize them as inhibitors of prostaglandin E2 production with the ability to produce anti-inflammatory effects that are approximately thirty times more potent than acetylsalicylic acid, better known as aspirin (Barrett et al., 1985; 1986). However, a broader understanding of cannflavins’ influence on cell biology in health and disease did not progress much since their initial description because of challenges associated with their extraction and the various political landscapes that limited their distribution. Nevertheless, some pre-clinical studies have provided intriguing new details about these molecules in recent years. For instance, the unnatural isomer of cannflavin B named FBL-03G (also known as caflanone) was found to increase apoptosis of pancreatic cancer cells *in vitro* while animal tests showed that the same small molecule could limit progression of metastatic pancreatic cancer (Moreau et al., 2019). Additionally, another study reported a possible neuroprotective effect of cannflavin A at concentrations lower than 10 µM that was attributed to the ability of this molecule to reduce aggregation of β amyloid through direct docking in the hydrophobic groove of the protein (Eggers et al., 2019). While these different findings provide novel insights about the pharmacological potential of cannflavins, the full range of molecular changes induced by cannflavins in cells remains to be described. To address this gap in our understanding of cannflavin pharmacology, we therefore focused on identifying novel mechanisms of action of these two related cannabis-derived metabolites in neuronal cells.

**Figure 1.**
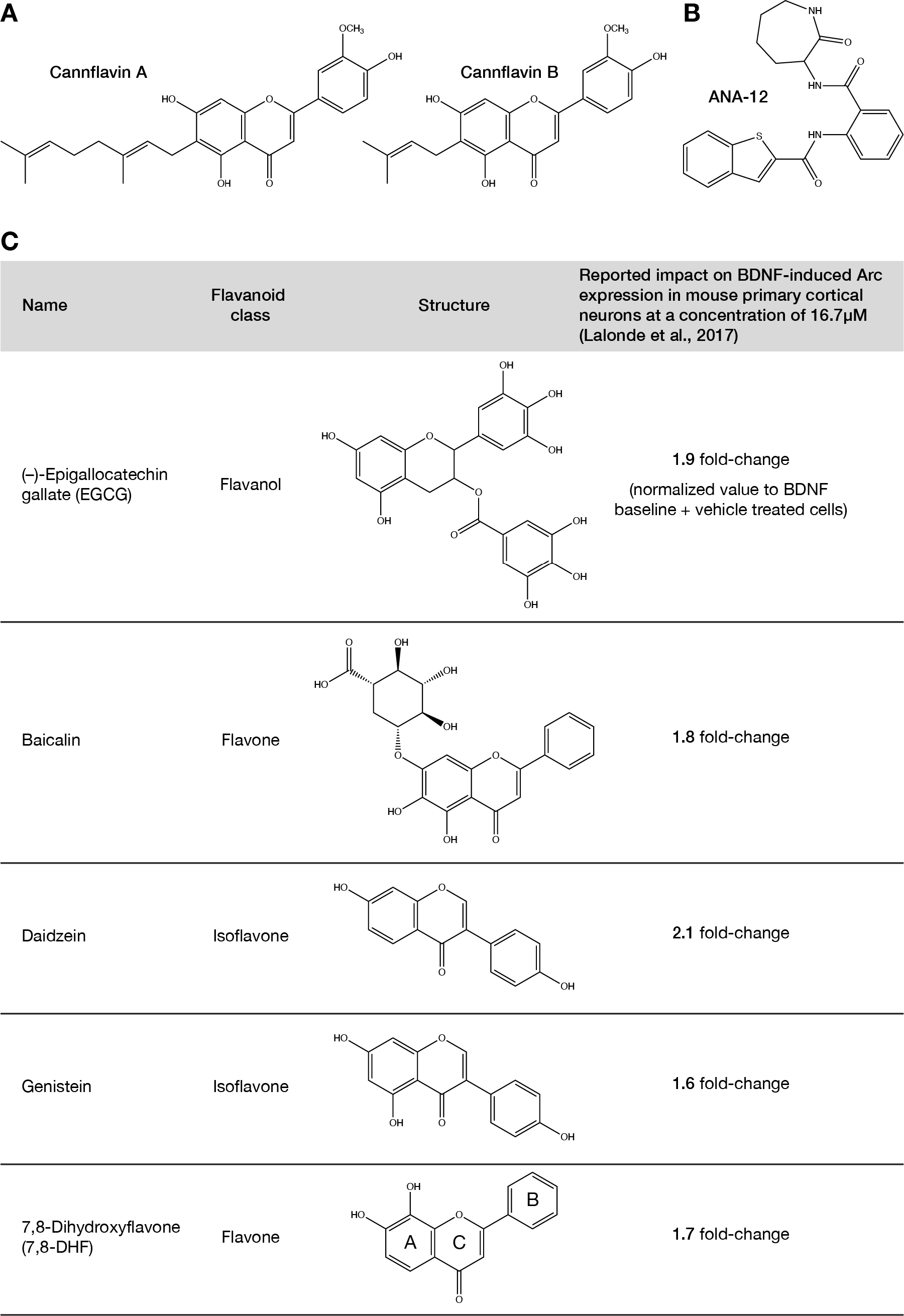
Chemical structures of key compounds. **A)** Chemical structure for cannflavin A and cannflavin B. **B)** Structure for the small molecule TrkB inhibitor ANA-12. **C)** Table presenting flavonoids organized by name, flavonoid class, chemical structure, and impact of BDNF-induced Arc expression in mouse primary cortical neurons as reported in Lalonde et al. (2017). Position of each ring is labeled for 7,8-dihydroxyflavone.

Cannflavins A and B are prenylated and highly lipophilic small molecules (Barrett et al., 1985; Choi et al., 2004), a characteristic that allows them to easily accumulate into cells where they can then presumably interact with different membrane-bound enzymes and receptors. Previously, we published a chemogenomic analysis that aimed at identifying small molecule modulators of Activity-regulated cytoskeleton-associated protein (Arc), which is a key regulator of neuroplasticity and cognitive functions (Bramham et al., 2010; Korb et al., 2011; Kedrov et al., 2019). Our approach in that project exploited the ability of the growth factor brain-derived neurotrophic factor (BDNF) to promote abundant *Arc* mRNA expression followed by nuclear accumulation of the protein product in mouse primary cortical neurons via activation of Tropomyosin receptor kinase B (TrkB) receptor (Lalonde et al., 2017). Here, we have adapted this assay to test the two cannflavins and found evidence of TrkB signaling interference by both molecules. These results then led us to complete a secondary high-throughput screen to test possible agonist activity of these flavonoids on G protein-coupled receptors (GPCRs), as well as other biochemical assays to confirm the influence of cannflavins and pinpoint their potential target engagement. These specific efforts suggest a model where cannflavins interfere with TrkB activity through direct inhibitory action on the receptor. Finally, image-based cellular test with immortalized Neuro2a cells ectopically expressing TrkB allowed us to demonstrate the capacity of cannflavins to block BDNF-induced neurite outgrowth. In summary, our study supports the classification of cannflavins as candidate inhibitors of TrkB receptor signaling in neuronal cells.

## 2. METHODS

### 2.1 Cell culture and transfection

Developing cerebral cortex from E16.5 CD-1 mouse embryos were dissected and then dissociated in trypsin solution for 15 min followed by three washes with phosphate-buffered saline (PBS). Trypsinized tissue was gently triturated to produce single cell suspension. Next, cells were seeded in poly-L-lysine/laminin coated 6-well plates at a density of 1.5 × 10^6^ per well and maintained in Neurobasal medium containing B27 supplement (2%, Invitrogen, Grand Island, NY), penicillin (50 U/ml, Invitrogen), streptomycin (50 µg/ml, Invitrogen) and glutamine (1 mM, Sigma). For experiments involving BDNF (PeproTech, Rocky Hill, NJ), the growth factor was added directly to the culture medium at a final concentration of 100 ng/ml for the indicated period of time.

Preparation of mouse primary cortical neuron cultures was approved by the University of Guelph Animal Care Committee and carried out according to institutional guidelines.

For neurite outgrowth assay, Neuro2a cells were cultured in DMEM [supplemented with 10% HyClone FetalClone II serum (Cytiva, Global Life Sciences Solutions, Marlborough, MA), penicillin (50 units/ml), and streptomycin (50 µg/ml)] and transfected overnight using Lipofectamine 2000 (Invitrogen) according to the manufacturer’s protocol.

### 2.2 Antibodies, plasmid, and pharmacological compounds

The anti-Arc rabbit polyclonal affinity purified antibody (#156 003) was purchased from Synaptic Systems (Goettingen, Germany). The antibody recognizing p42 Mapk (Erk2, sc-1647) was from Santa Cruz Biotechnology (Santa Cruz, CA). The antibodies recognizing phosphorylated TrkA^Tyr490^/TrkB^Tyr516^ (#4619), phosphorylated p44/42 Mapk (Erk1/2^Thr202/Tyr204^, #4370), Akt (#4691), phosphorylated Akt^Thr308^ (#2965), phosphorylated Akt^Ser473^ (#4060), mTor (#2983), phosphorylated mTor^Ser2448^ (#2971), and phosphorylated rpS6^Ser240/244^ (#2215) were acquired from Cell Signaling Technology (Beverly, MA). The antibody detecting TrkB (MAB397) was acquired from R&D Systems (Minneapolis, MN), and the one specific for mouse phosphorylated TrkB^Tyr705^ (ab229908) was from Abcam (Waltham, MA). The β-actin (A1978) and M2 FLAG (F1804) antibodies were from Sigma-Aldrich (St-Louis, MO), while the Map2 (AB5543) antibody was purchased from EMD Millipore Corps (Billerica, MA). Finally, cross-absorbed horseradish peroxidase-conjugated secondary antibodies were from Invitrogen.

The pCMV6-Ntrk2-Myc-DDK (FLAG) plasmid (MR226130) was purchased from OriGene Biotechnologies (Rockville, MD). ECGC, genistein, and daidzein were from Sigma-Aldrich.

ANA-12 (Figure 1B) was from Tocris Bioscience (Bristol, UK) and U0126 was from Biosciences (Thermo Fisher).

The synthesis and purification of cannflavins A and B were produced using the method of Rea and colleagues (2019). Briefly, *Cannabis sativa* L. prenyltransferase 3 (CsPT3) was recombinantly expressed in *Saccharomyces cerevisiae* and the microsomal fraction containing CsPT3 was collected for *in vitro* enzyme assays. Assays containing 200 µM chrysoeriol, 400 µM GPP or DMAPP, 1 mg/mL of microsomal protein, and 10 mM MgCl2 in 100 mM Tris-HCl buffer were conducted at 37°C for 120 min and terminated with the addition of 20% formic acid. Cannflavin products were extracted with three volumes of ethyl acetate, the organic layer was dried under N2 gas and resuspended in methanol. The products were purified by high-performance liquid chromatography (HPLC) on an Agilent 1260 Infinity system equipped with a Waters SPHERISORB 5 µm ODS2 column, and eluted with a 20 min linear gradient from 45% to 95% methanol in water containing 0.1% formic acid. Product identities were confirmed via liquid chromatography-mass spectrometry (LC-MS) according to published methods (Rea et al., 2019) (Supplementary Figure 1). Cannflavins produced *in vitro* were dried under nitrogen gas and resuspended in dimethyl sulfoxide (DMSO). The final products were confirmed via HPLC, as described above, and quantified by absorption at 340 nm relative to authentic standards.

### 2.3 Western blotting

For western blot analyses, cells were collected by scraping in ice-cold radioimmunoprecipitation assay (RIPA) buffer (50 mM tris-HCl [pH 8.0], 300 mM NaCl, 0.5% Igepal-630, 0.5% deoxycholic acid, 0.1% SDS, 1 mM EDTA) supplemented with a cocktail of protease inhibitors (Complete Protease Inhibitor without EDTA, Roche Applied Science, Indianapolis, IN) and phosphatase inhibitors (Phosphatase Inhibitor Cocktail 3, Sigma-Aldrich). One volume of 2× Laemmli buffer (100 mM tris-HCl [pH 6.8], 4% SDS, 0.15% bromophenol blue, 20% glycerol, 200 mM β-mercaptoethanol) was added and the extracts were boiled for 5 min. Samples were adjusted to an equal concentration after protein concentrations were determined using the BCA assay (Pierce, Thermo Fisher Scientific). Lysates were separated using SDS–PAGE (polyacrylamide gel electrophoresis) and transferred to a nitrocellulose membrane. After transfer, the membrane was blocked in TBST (tris-buffered saline and 0.1% Tween 20) supplemented with 5% nonfat powdered milk and probed with the indicated primary antibody at 4°C overnight. After washing with TBST, the membrane was incubated with the appropriate secondary antibody and visualized using enhanced chemiluminescence (ECL) reagents according to the manufacturer’s guidelines (Pierce, Thermo Fisher Scientific).

The following procedure was used to quantify western blot analyses. First, equal quantity of protein lysate as determined by the BCA assay was analyzed by SDS-PAGE for each biological replicate. Second, the exposure time of the film to the ECL chemiluminescence was the same for each biological replicate. Third, all the exposed films were scanned on a HP Laser Jet Pro M377dw scanner in grayscale at a resolution of 300 dpi. Fourth, the look-up table (LUT) of the scanned tiff images was inverted and the intensity of each band was individually estimated using the selection tool and the histogram function in Adobe Photoshop CC 2021 software. Finally, the intensity of each band was divided by the intensity of its respective loading control (β-actin) to provide the normalized value used for statistical analysis.

### 2.4 Immunocytochemistry

Indirect immunofluorescence detection of antigens was carried out using cortical neurons cultured on poly-L-lysine/laminin coated coverslips in 24-well plates at a density of 0.1 × 10^6^ per well. After experimental treatment, cells were washed twice with phosphate-buffered saline (PBS) and fixed for 30 min at room temperature with 4% paraformaldehyde in PBS. After fixation, cells were washed twice with PBS, permeabilized with PBST (PBS and 0.25% Triton X-100) for 20 min, blocked in blocking solution (5% goat non-immune serum in PBS) for another 30 min, and finally incubated overnight at 4°C with the first primary antibody in blocking solution. The next day, coverslips were extensively washed with PBS and incubated for 2 hours at room temperature in the appropriate fluorophore-conjugated secondary antibody solution [Alexa Fluor 488-, Alexa Fluor 594, or Alexa Fluor 647-conjugated secondary antibody (Molecular Probes, Invitrogen) in blocking solution]. After washes with PBS, the coverslips were incubated again overnight in primary antibody solution for the second antigen, and the procedure for conjugation of the fluorophore-conjugated secondary antibody was repeated as above. Finally, cell nuclei were counterstained with 4′,6-diamidino-2-phenylindole (DAPI), and coverslips were mounted on glass slides with ProLong Antifade reagent (Invitrogen).

Cells cultured on coverslips from three independent biological replicates were imaged with a Nikon Eclipse Ti2-E inverted microscope equipped with a motorized stage, image stitching capability, and a 60× objective (Nikon Instruments, Melville, NY). A total of 30 images per condition were analyzed, three biological replicates were performed where ten images were analyzed at random. All cells within the 60X field were included in the analysis. Image analysis was performed with ImageJ and NIS Elements and the following procedure was used to quantify nuclear Arc level in response to BDNF-TrkB signaling. First, original raw tiff files were opened and the nucleus of all neurons in the image was located based on Map2 or NeuN immunostaining, then average pixel intensity corresponding to Arc immunofluorescence was measured for a 30- pixel spot positioned at the center of the nuclear compartment. Second, for each measure of Arc nuclear immunofluorescence pixel intensity, a measure of background pixel intensity from the same image channel was acquired and subsequently subtracted from the Arc nuclear immunofluorescence pixel intensity value. Finally, Arc immunofluorescence signal from untreated samples was used to establish an objective threshold (two standard deviations above the nuclear Arc immunofluorescence signal averaged from a representative population of untreated neurons) which allowed for comparison of nuclear Arc expression between different experimental conditions.

### 2.5 Real-time reverse transcriptase PCR

After experimental treatment, total RNA was isolated from primary cortical neuron cultures using the TRIzol method (Invitrogen). The concentration of total RNA was measured using a NanoDrop ND-8000 spectrophotometer (Thermo Fisher Scientific) and first-strand complementary DNA (cDNA) was synthesized using the iScript cDNA Synthesis Kit (Bio-Rad, Hercules, CA). Real-time PCRs were performed using gene-specific primers and monitored by quantification of SYBR Green I fluorescence using a Bio-Rad CFX96 Real-Time Detection System. Expression was normalized against *Gapdh* expression. The relative quantification from three biological replicates was calculated using the comparative cycle threshold (ΔΔCT) method.

Primers for real-time reverse transcription PCR experiments were: *Arc* primer pair one, 5′-

TAGCCAGTGACAGGACCCAG-3′ (forward) and 5′-CAGCTCAAGTCCTAGTTGGCAAA-3′ (reverse);

*Arc* primer pair two, 5′- CGCCAAACCCAATGTGATCCT-3′ (forward) and 5′-

TTGGACACTTCGGTCAACAGA-3′ (reverse); *Gapdh*, 5′-ATGACCACAGTCCATGCCATC-3′ (forward) and 5′-CCAGTGGATGCAGGGATGATGTTC-3′ (reverse).

### 2.6 PRESTO-Tango GPCR assay

Parallel receptorome expression and screening via transcriptional output, with transcription activation following arresting translocation (PRESTO-Tango) was used to assess cannflavin A and cannflavin B potential to stimulate G protein-coupled receptors (GPCRs) according to published methods (Kroeze et al., 2015). Overall, 320 distinct nonolfactory human GPCRs were tested.

### 2.7 Neurite outgrowth assessment

Neuro2A cells transfected with a pCMV6-Ntrk2-Myc-DDK (FLAG) construct were selected with G-418 (Geneticin) to produce a stable cell line that constitutively expresses Myc-FLAG tagged- TrkB. For neurite outgrowth assessment, cells were seeded on 15 mm glass coverslips in 12-well plates at a density of 2.0 × 10^4^ per well and allowed to attach overnight. Next day, cells were treated with recombinant BDNF (1 nM) plus cannflavins (10 µM), ANA-12 (10 µM), or vehicle control (DMSO). Phase contrast digital images were collected with a Nikon Eclipse Ti2-E inverted microscope and a 20× objective 24 hours after start of treatment (five fields per dish, three wells per condition). Image analysis was completed using ImageJ software using the following procedure. First, any cells with a fragmented nucleus were excluded from the analysis, therefore the total number of viable cells was counted per field. Second, for all identified viable cells the total number of neurites and number of cells with neurites longer than 2 cell bodies in diameter were counted per field.

### 2.8 Alamar Blue assay

Neuro2A cells stably expressing pCMV6-Ntrk2-Myc- DDK (FLAG) were seeded in 96-well plates at a density of 1.0 x 104 and allowed to attach overnight. Next day, cells were treated with recombinant BDNF (1 nM) plus cannflavins (10 µM), ANA-12 (10 µM), or vehicle control (DMSO). Resazurin sodium salt (Sigma-Aldrich) was dissolved in Hank’s balanced salt solution (Gibco) to make a 0.5 mM working solution. After 21 hours of treatment, working solution was diluted 1:10 into the 96-well plate and incubated at 37°C for 3 hours. Fluorescence was measured with excitation/emission at 570/590 nm using a fluorescence microplate reader. All samples were completed in triplicate, and a background control reading (media with working solution) was subtracted from each value.

### 2.9 Statistics

Unless mentioned otherwise, all results represent the mean ±SEM from at least three independent experiments. ANOVA followed by Tukey’s or Dunnett’s post hoc test for multiple comparisons were performed where indicated.

## 3. RESULTS

### 3.1 Impact of cannflavins on BDNF-induced Arc expression in mouse primary cortical neurons

Previously, we completed a chemogenomic screen with primary mouse cortical neurons that identified a suite of compounds that acted as Arc expression modifiers (Lalonde et al., 2017). Part of this group included five distinct flavonoids (Figure 1C)—namely (–)-epigallocatechin (ECGC), baicalin (BAI), 7,8-dihydroxyflavone (7,8-DHF), daidzein, and genistein—which were found to enhance nuclear Arc protein levels above the control measure when co-applied at a final concentration of 16.7 µM with recombinant BDNF for 6 h. Searching for a possible explanation to this phenomenon, we were intrigued by several studies that had linked each of these five flavonoids to either enhancement of *BDNF* and/or *TrkB* mRNA expression, or to the potentiation of downstream TrkB-dependent signaling (Pan et al., 2012; Gundimeda et al., 2014; Ding et al., 2018; Lu et al., 2019). Based on these observations, we hypothesized that cannflavin A and cannflavin B could act in a similar fashion and promote Arc protein abundance when added to cultured mouse cortical neurons that were stimulated with exogenous BDNF. Unexpectedly, though, western blot analysis assessing BDNF-induced Arc expression in conjunction with cannflavins with concentrations of the flavonoids ranging between 1 to 20 µM revealed an opposite effect. Specifically, we found that application of cannflavins to cell culture media prevented the normal increase in Arc protein by BDNF in a dose-dependent manner where 10-20 µM of cannflavin A, and all tested concentrations (1-20 µM) of cannflavin B, resulted in significantly less Arc protein abundance than seen in the BDNF-alone control measure (Figure 2A). To further support this result, we repeated the experiment using fluorescent immunocytochemistry and quantified nuclear Arc changes, as we had done previously in our chemogenomic screen (Lalonde et al., 2017). Of note, we also tested the flavonol ECGC, as well as the isoflavone daidzein and genistein, since those molecules were found to produce the opposite effect of increasing BDNF- induced nuclear Arc levels according to our previous screening results. As shown in Figure 2B and Supplementary Figure 2, cells that were co-treated with BDNF and 10 µM daidzein or genistein presented a moderate increase in the percentage of nuclei with Arc expression above threshold in comparison to the BDNF-alone control. Interestingly, though, the ECGC condition was similar on average to the BDNF-alone condition suggesting that the final concentration of this specific flavonoid must be greater than 10 µM to have an impact of TrkB-induced nuclear Arc level. Most importantly, and consistent with our western blot analysis above, cultures treated with BDNF and 1-10 µM cannflavins presented similar overall trends in the reduction of Arc-positive neuronal abundance in comparison to the unstimulated control (Figure 2C). Together, these results confirm the observed discrepancy between cannflavins and the other flavonoids found in our earlier screen, which focused on BDNF-induced Arc expression modifiers.

**Figure 2.**
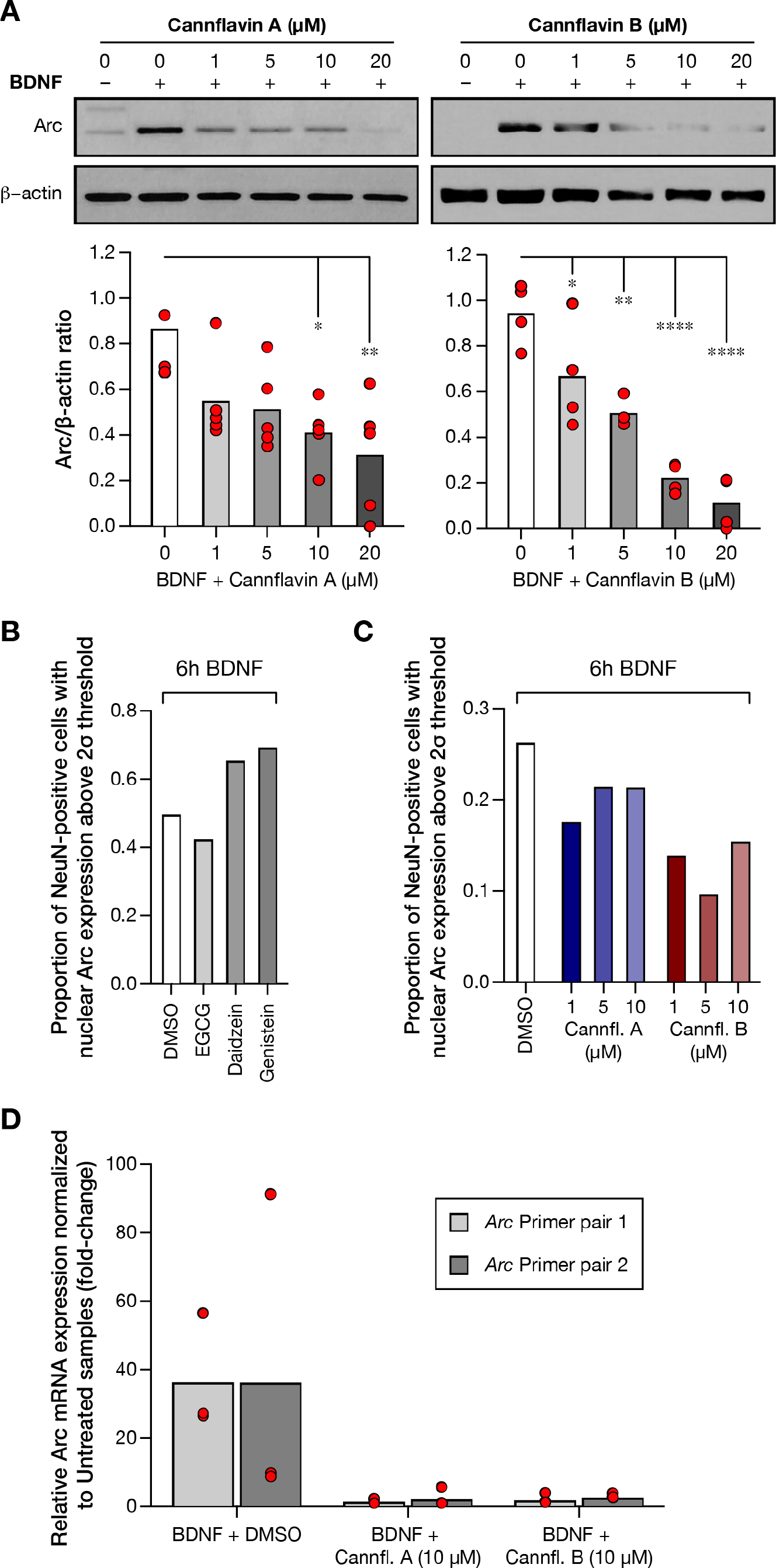
Cannflavin A and cannflavin B decrease BDNF-induced Arc protein and mRNA levels in mouse primary cortical neurons. **A)** Western blot and corresponding densitometry analysis showing Arc protein abundance in mouse primary cortical cultures when treated with BDNF and various concentrations (0, 1, 5, 10, 20 µM) of cannflavin A (left) or cannflavin B (right). β-actin was used a loading control and graphs show mean of Arc/β-actin ratio for each condition. Biological replicates: *n =* 5 (cannflavin A), *n =* 4 (cannflavin B). One way ANOVA revealed a significant decrease in the abundance of Arc with increasing cannflavin concentrations. Cannflavin A (*F*4,20 = 4.568, *p* = 0.0088), Tukey’s post-hoc test, * *p <* 0.05, ** *p* < 0.001. Cannflavin B (*F*4,15 = 24.07, *p* < 0.0001), Tukey’s post-hoc test, * *p <* 0.05, ** *p* < 0.001, **** *p* < 0.0001. **B)** Quantification of immunocytochemistry coverslips treated with various flavonoids (EGCG, daidzein, and genistein; all 10 µM final concentration). Quantification was completed by using a ratio of Map2-positive cells with nuclear Arc above the 2σ nuclear Arc pixel intensity in the control condition (BDNF treatment alone). **C)** Quantification of immunocytochemistry coverslips treated with various concentrations (0, 1, 5, 10 µM) of cannflavin A (blue bars) or cannflavin B (red bars). Quantification was completed similar to B. **D)** Quantitative real-time PCR *Arc* mRNA analysis using two Arc primer pairs shows a decrease in *Arc* transcripts when treated with 10 µM of cannflavin A or cannflavin B, relative to cells treated with BDNF alone.

Next, to ascertain whether the cannflavins’ influence on TrkB signaling and Arc protein levels was produced before or after gene transcription, we performed a quantitative polymerase chain reaction (qPCR) experiment using two pairs of primers targeting different regions of the *Arc* transcript. Here, comparison of *Arc* mRNA abundance between untreated, BDNF-alone control, and BDNF with cannflavin A (10 µM) or cannflavin B (10 µM) samples clearly indicated that cannflavins prevent induction of *Arc* mRNA expression (Figure 2D), therefore suggesting that the effect of these compounds must occur somewhere between the activation of TrkB receptors by BDNF and the activation of the transcriptional machinery involved in *Arc* expression.

### 3.2 Evaluating agonist potential of cannflavins on Tango GPCR assay

Considering the transactivation crosstalk between GPCRs and receptor tyrosine kinases, including TrkB (Rajagopal et al., 2004; 2006; El Zein et al., 2007), and recent evidence for GPCR modulation/self-association by flavonoids on the latter (Herrera-Hernández et al., 2017; Ortega et al., 2019), we speculated that one mechanism by which cannflavins could interfere on *Arc* mRNA expression in cortical neurons involves activation of a G protein signal that transinactivates the function of molecular cascades downstream of TrkB receptors responsible for *Arc* expression. To explore this scenario, we tested the effect of cannflavins on the GPCRome, *en masse,* using the PRESTO-Tango assay—an unbiased high-throughput screening approach adapted to identify agonist activity of agents towards the large family of GPCRs (Kroeze et al, 2015). Interestingly, applying cannflavin A revealed no effect on any of the 320 different GPCRs tested while cannflavin B was found to produce only weak increase (4.4 fold-change) of GPR150 activity from baseline, a negligible effect in comparison to the positive control (51.3 fold-change, dopamine D2 receptor stimulated by quinpirole) (Figure 3). Faced with these results, we then re-focused our attention on the possibility that cannflavins act more directly on the TrkB receptor and/or its downstream signaling components.

**Figure 3.**
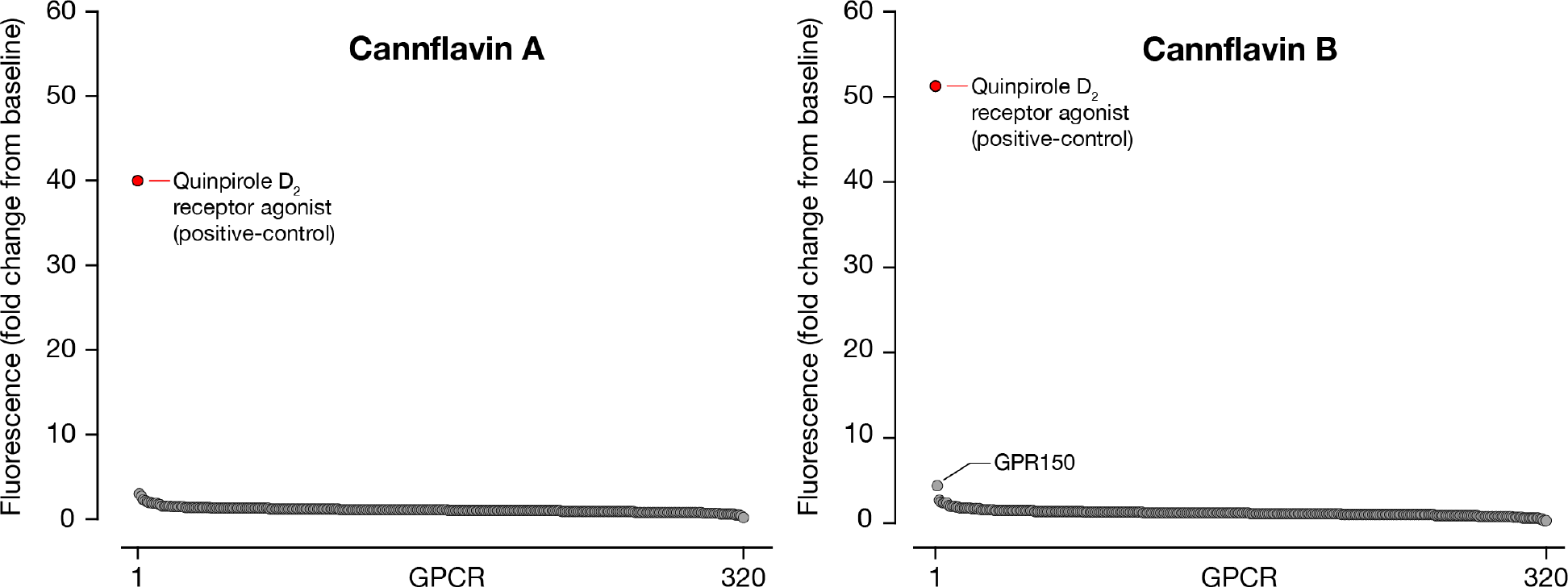
Cannflavins do not stimulate G protein-coupled receptors signaling. Data for 320 GPCRs are presented as an average fold change from baseline upon compound addition. Application of quinpirole to D2 receptor is used as positive control in each plate. Compounds were used at 10 μM and all tests run in quadruplicate.

### 3.3 Elucidating cannflavins effects on TrkB signaling

BDNF binding to the extracellular domain of a TrkB receptor stimulates its dimerization and the phosphorylation of various intracellular tyrosine residues which is followed by the recruitment of pleckstrin homology (PH) and SH2 domain containing proteins—such as FRS2, Shc, SH2B, and SH2B2—that regulate distinct concurrent signaling cascades (Qian et al., 1998; Meakin et al., 1999). To explore whether cannflavins interfere with the activation of TrkB receptors by BDNF in primary cortical neurons, we used a western blotting approach and probed lysates with two P- Trk antibodies: one specific to mouse P-TrkA/B^Y499/515^, which is associated with the recruitment of Shc adaptor proteins, and another specific to mouse P-TrkB^Y705^ in the catalytic domain of the receptor (Figure 4A). This approach revealed that, indeed, cannflavins can prevent BDNF from effectively stimulating its target receptor (Figure 4B). To further support this result, we tested the activation of signaling pathways that are likely regulated downstream of TrkB, including the Ras-Raf-MEK-Mapk and the PI3K/Akt/mTor cascades (Huang et al., 2003; Kowiański et al., 2018) (Figure 4A). Interestingly, our analyses revealed that both cannflavin A and cannflavin B sharply reduced normal increase in P-Mapk, P-Akt, P-mTor, and P-rpS6 levels produced by BDNF (Figure 4C). The fact that there were those changes in these three molecular pathways, which to a great extent occurs in parallel with limited cross-interaction (Kowiański et al., 2018), strongly suggest that cannflavins must act at an early stage in TrkB signal activation.

**Figure 4.**
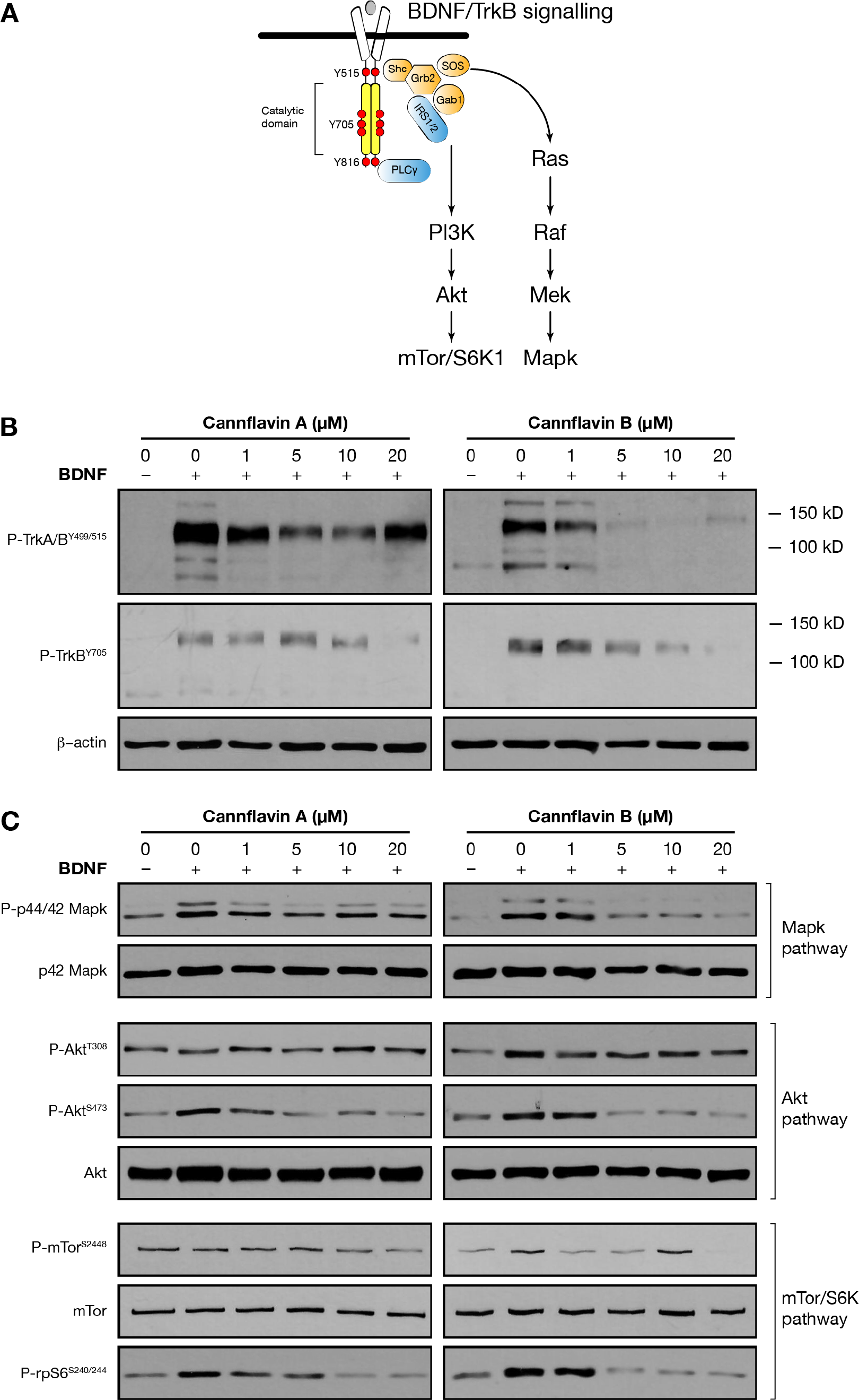
Cannflavins A and B decrease activation of downstream pathways of TrkB. A) Simplified schematic of BDNF activation of TrkB receptors and downstream signaling pathways. **B)** Western blot analysis showing phosphorylation of TrkB when treated with BDNF and various concentrations (0, 1, 5, 10, 20 µM) of cannflavin A (left) or cannflavin B (right). β-actin was used a loading control. **C)** Western blot analysis showing phosphorylation of Mapk, Akt, and mTor proteins when treated with various concentrations (0, 1, 5, 10, 20 µM) of cannflavin A (left) or cannflavin B (right).

### 3.4 Functional characterization TrkB inhibition by cannflavins on BDNF-dependent neurite outgrowth

Our biochemical analyses with mouse primary cortical neurons suggest that cannflavins A and B have inhibitory activity towards TrkB receptors. In order to establish if this effect is sufficient to limit cellular processes under the control of BDNF signaling, we used neuroblastoma Neuro2a cells stably expressing Ntrk2 (TrkB)-Myc-FLAG to complete a neurite outgrowth experiment (Figure 5A). As shown in Figure 5B, Neuro2a cells have low TrkB expression with negligeable phosphorylation of the receptor under basal conditions, while cells stably expressing the receptor display greater responsiveness to exogenous application of BDNF—which is clearly demonstrated by higher level of P-TrkB. Further, pre-application of cannflavins (20 µM) with BDNF to the culture media for 6 h reduced BDNF-induced TrkB phosphorylation (Figure 5C). Interestingly, we did not observe a decrease in P-Mapk levels with treatment of ANA-12 or cannflavins as was observed in cortical neurons (Figure 4C), a distinction that we attribute to the fact that overexpression of TrkB in Neuro2A produced maximal phosphorylation of the p42 subunit of Mapk and no change in signal when BDNF was added to the Neuro2a cells (Figure 5B). Most importantly, neurite assays completed with and without the application of ANA-12, cannflavin A, and cannflavin B revealed that all three compounds produced a significant decrease in the total number of neurites per field (Figure 5D and E) and number of cells with processes twice the length of the cell (Figure 5F) when applied at a final concentration of 10 µM to culture media. Cell viability was then measured using Alamar Blue cell survival assay, which revealed no significant difference between the BDNF control condition and the BDNF plus ANA-12 or cannflavins (Figure 5G). Of note, our measures of total number and length of neurites for the ANA-12 and cannflavins conditions in this experiment were comparable to the baseline values observed in wild-type Neuro2a cells not overexpressing TrkB (Supplementary Figure 3). Altogether, our results here strongly support that cannflavins act on TrkB receptors, preventing BDNF activation of downstream signaling of the receptor.

**Figure 5.**
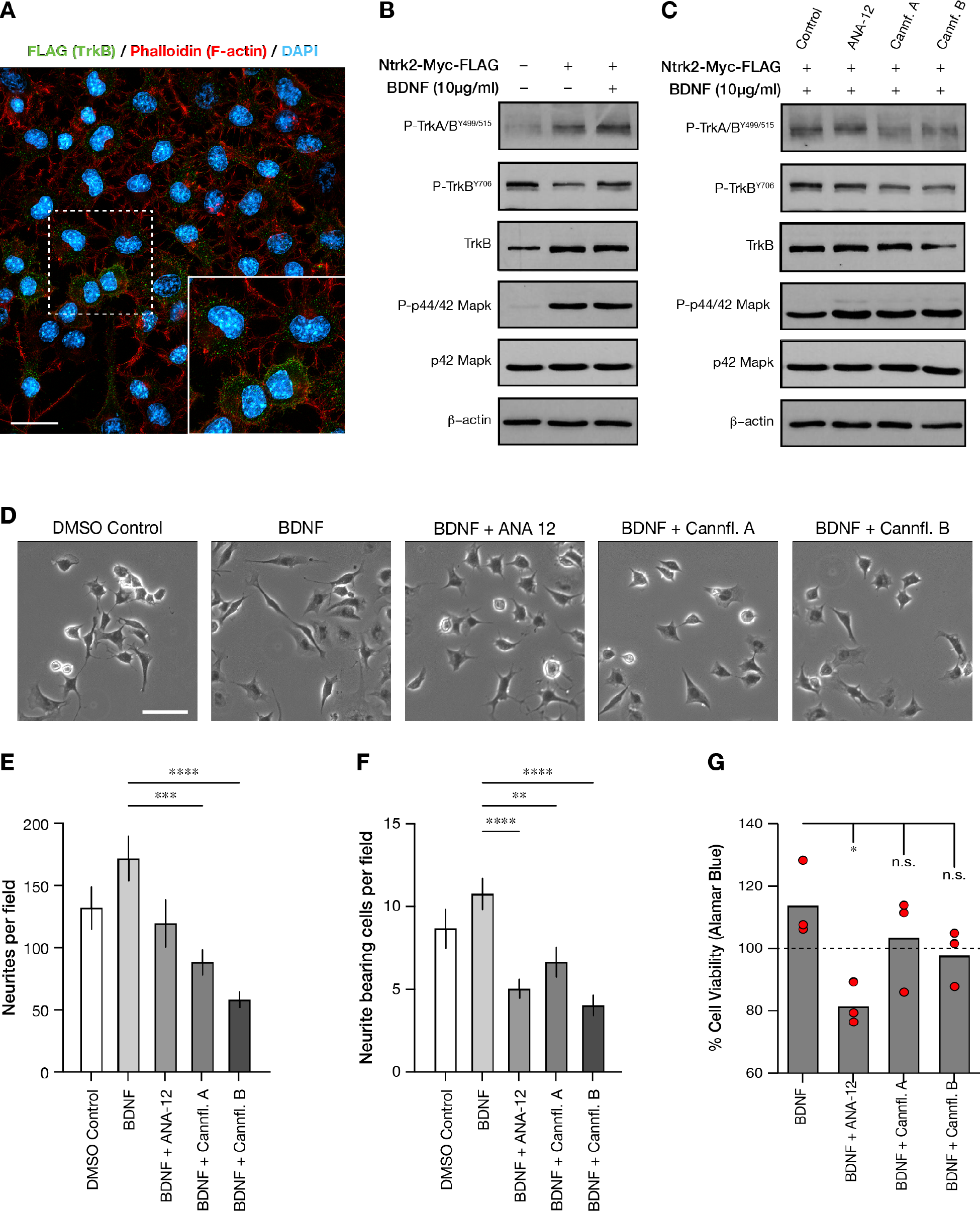
Cannflavin A and cannflavin B reduces BDNF-induced neurite outgrowths in neuroblastoma cells. **A)** Neuro2a cells were transfected to stably express Ntrk2-Myc-FLAG. Immunocytochemistry was completed to validate that the cells were successfully transfected. Scale bar = 20 µM. **B)** Western blot analysis showing phosphorylation of TrkB and Mapk in regular Neuro2a cells compared to those stably expressing Ntrk2-Myc-FLAG. β-actin was used a loading control. **C)** Western blot analysis showing phosphorylation of TrkB and Mapk when treated with 10 µM of ANA-12, cannflavin A, or cannflavin B with the addition of BDNF (10 µg/mL). The small-molecule non-competitive TrkB antagonist ANA-12 was used as a positive control (Cazorla et al., 2011) and β-actin was used as a loading control. **D)** Phase contrast images of Ntrk2-Myc- FLAG Neuro2as treated with or without BDNF (10 µg/mL) and 10 µM of ANA-12, cannflavin A, or cannflavin B. Scale bar = 50 µm. Images were quantified by counting **E)** total number of neurites, and **F)** total number of cells bearing neurites twice the length of cell body. One-way ANOVA was used to analyze data. Total number of neurites (*F*4,150 = 8.264, *p* < 0.001); total number of cells bearing neurites (*F*4,150 = 9.433, *p* < 0.001); Dunnett’s multiple comparisons test to BDNF condition, * *p* < 0.05; ** *p* < 0.01; *** *p* < 0.001; *** *p* < 0.0001. Graphs represent mean ±SEM. **G)** Cell viability was measured using an Alamar Blue reaction after 24 hours of treatment (BDNF + ANA-12, cannflavin A, or cannflavin B). Application of ANA-12 produced a significant decrease in cell viability compared to DMSO control but not cannflavin A or cannflavin B. One- way ANOVA was used to analyze data. Cell survival main effect (*F*4,10 = 3.991, *p* = 0.0345). Tukey’s multiple comparisons test, * *p* < 0.05. Of note, comparison between DMSO and ANA- 12, specifically, was not significant (*p* = 0.2532). Graphs represent mean (3 biological replicates) and dash line DMSO reference measure.

## 4. DISCUSSION

This study provides evidence for an inhibitory effect of cannflavins A and B, two flavonoids from *C. sativa*, on BDNF-induced Arc expression through disruption of TrkB receptor signaling. These results contrast our previous observations that other flavonoids exhibit a potentiating effect towards BDNF-induced Arc accumulation in mouse primary cortical neurons (Lalonde et al., 2017). Strikingly, the addition of cannflavin A or cannflavin B to the culture medium of mouse primary cortical neurons at a minimum concentration of 5 µM consistently prevented the induction of *Arc* mRNA expression, suggesting that these molecules act between BDNF-activation of TrkB receptors and transcription of the *Arc* gene. Therefore, we subsequently investigated the impact of cannflavins on the downstream pathways of TrkB using biochemical analyses and uncovered a consistent decrease in the activation of the Ras-Raf-Mek-Mapk and the PI3K/Akt/mTor cascades. In connection with these results, we demonstrated that cannflavins inhibited BDNF-induced neurite outgrowth in neuroblastoma Neuro2a cells stably overexpressing TrkB. Taken together, our study provides a new path to better understand the effects that have been reported in recent years about cannflavins and other closely related compounds against certain cancer cell types (Brunelli et al., 2009; Moreau et al., 2019).

### 4.1 Structural determinants of cannflavins activity towards TrkB

All flavonoids have a basic flavan nucleus with two aromatic rings (the A and the B rings) interconnected by a three-carbon-atom heterocyclic ring (the C ring), as illustrated in Figure 1C for the flavone 7,8-dihydroxyflavone (7,8-DHF, also known as tropoflavin). Interestingly, a previous study focusing on 7,8-DHF, which is a compound reported to mimic the physiological activity of BDNF and stimulate TrkB signaling *in vitro* and *in vivo* (Jang et al., 2010; Zeng et al., 2012), helps speculate about what structural feature could provide TrkB antagonistic activity to cannflavins. Specifically, previous comparison of different 7,8-DHF derivatives on TrkB phosphorylation and downstream Akt signaling revealed that the presence of a 3’-hydroxy group (or to a lesser extent a 2’-hydroxy group) on the B ring confers a TrkB stimulatory effect to a 7,8- DHF derivative compound whereas addition of a 4’-hydroxy group, as shown in Figure 6A with 7, 8, 4’-trihydroxyflavone and Figure 6B with 3, 5, 7, 8, 3, 4’-hexahydroxyflavone, inversely mediates inhibition of the receptor (Liu et al., 2010). Since cannflavins A and B are both hydroxylated at the 4’ position on the B ring (Figure 1A), similar to compounds found by Liu and colleagues (2010) interfering with TrkB phosphorylation in rat primary cortical neurons (Figures 6A and 6B), we thus suspect that this specific structural feature is key in mediating the TrkB antagonistic effect observed in our study. In this context, it will be revealing whether cannflavin derivatives with different patterns of hydroxylation on the B ring produce a different activity towards TrkB and downstream cellular effects.

**Figure 6.**
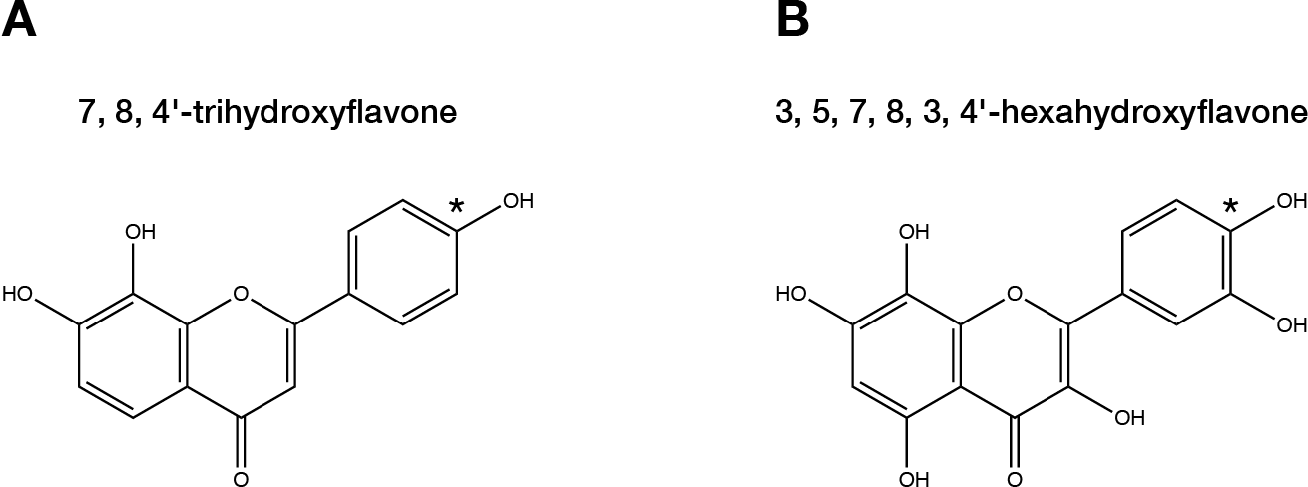
Chemical structure of 7,8-dihydroxyflavone derivatives reported to block TrkB signaling in Liu and colleagues (2010). Asterisk indicates hydroxylated 4’ position on B ring of each compound.

In addition to 4’ hydroxylation of the B ring, prenylation is another structural element that we must consider in relation to the difference we consistently observed between cannflavin A and cannflavin B towards TrkB. As seen in Figure 1A, cannflavin B has only one isoprene unit while cannflavin A has two, making the later an overall larger and more lipophilic molecule. Consequently, we speculate that the smaller size of cannflavin B may facilitate access and/or create a stronger binding affinity to TrkB in a cellular context, which could then explain why we have been measuring a more potent inhibitory response of the receptor phosphorylation and blunting of downstream signaling with this specific compound. Here, though, we acknowledge that one limitation to our study is we do not demonstrate direct interaction of cannflavins A and B with TrkB at this point. Nevertheless, two specific results from our study support to a certain degree the idea that a direct functional interaction most likely occur between cannflavins and TrkB. First, our interrogation of the GPCRome using the PRESTO-Tango assay showed that cannflavins do not stimulate the activity of more than 300 GPCRs, which rules out the possibility that cannflavins are disrupting TrkB function through a GPCR transinactivation event in our cellular experiments (Rajagopal et al., 2004; 2006; El Zein et al., 2007). And second, our experiment with Neuro2a cells stably expressing TrkB reveals that cannflavins block the growth of neurites, as well as the observed cell survival stimulatory effect, produced by exogenous BDNF application in this model. Since these phenotypes are directly tied to the overexpression of TrkB, and that application of cannflavins consistently returns neurite and survival measures to those of wild-type Neuro2a cells (Supplementary Figure 3), we consider these results as evidence for a direct interaction between cannflavins and TrkB. To conclude on this point, whether the activity of receptor tyrosine kinases other than TrkB is interfered by cannflavins remains unknown and should be considered in future research.

### 4.2 Therapeutic potential of cannflavins

Although cannflavins are recognized to produce potent anti-inflammatory effects by inhibiting the biosynthesis of various pro-inflammatory mediators, including microsomal prostaglandin E2 synthase-1 (mPGES-1) and 5-lipoxygenase (5-LO) (Barrett et al., 1985; 1986; Werz et al., 2014), our study cautions that these flavonoids may not be adequate to intervene against neuroinflammation or provide pro-cognitive effects because of their impact on TrkB signaling (Jaeger et al., 2018). Also, recent evidence suggests that the therapeutic action on mood of both typical and fast-acting antidepressants may be linked by their ability to directly bind TrkB receptors and allosterically increase BDNF signaling (Casarotto et al., 2021). Based on that specific information, and our results in this study, we consequently suspect that cannflavins may exacerbate depressive mood. Keeping those important details in mind, we nevertheless believe that cannflavin A and/or cannflavin B should be further studied in relation to clinical circumstances where overactive or dysregulated TrkB signaling is a contributing factor. For instance, increased TrkB expression was detected in low-grade astrocytoma and glioblastoma (Wadhwa et al., 2003; Assimakopoulou et al., 2007), while BDNF-induced activation of TrkB has been found to increase the viability of brain-tumor stem cells isolated from glioblastoma (Lawn et al., 2015). Furthermore, a study uncovered a link between the ability of glioblastoma to make less invasive cancer cells around them more aggressive via the transfer of TrkB-containing exosomes, revealing this way a mechanism by which these tumours can influence their environment to promote disease aggressiveness (Pinet et al., 2016), and other reports have shown that inhibition of TrkB-associated signaling may be an effective strategy to limit the formation of astrocytomas (Ni et al., 2017) as well as the survival of glioblastoma cancer cells (Pinheiro et al., 2017). Finally, accumulating evidence support that TrkB signaling could also be a therapeutic target for other cancer types, including lung (Sinkevicius et al., 2014; Chen et al., 2016), breast (Choy et al., 2017; Contreras-Zárate et al., 2019), and pancreatic cancer (Oyama et al., 2021). Taken together, these studies suggest that targeting TrkB signaling with cannflavins could provide therapeutic benefits against different cancer types, in particular those affecting the nervous system.

Lastly, peripheral increase in BDNF appears to be common in autism spectrum disorder based on meta-analysis evidence (Zheng et al., 2016), and the BDNF/TrkB pathway has been linked to hyperexcitability caused by axonal transection and some forms of epileptogenesis (Lin et al., 2020). Whether cannflavins could be valuable agents in those contexts should be tested. In conclusion, our study expands the range of cellular effect for cannflavins beyond inflammation and supports the examination in more detail of these compounds to attack brain cancer cells as well as mitigate aberrant neurodevelopmental and circuit phenotypes that could be resulting from uncontrolled BDNF-TrkB signaling.

## DATA AVAILABILITY STATEMENT

The raw data supporting the conclusions of this article will be made available by the authors, without undue reservation.

## ETHICS STATEMENT

The use of mice for primary neuron cultures was reviewed and approved by the University of Guelph Animal Care Committee.

## AUTHOR CONTRIBUTIONS

J.H., A.W.M., B.A., J.Y.K., and J.L. planned and designed the experiments. C.P., J.Y.K., and

T.A.A. provided key resources, and J.H., A.W.M., B.A., and C.A. performed the research. J.H., A.W.M., B.A., C.P., C.A., and J.L. analyzed the data and prepared figures. J.H., A.W.M., and J.L. wrote the manuscript with revisions from all other authors.

## ACKNOWLEDGMENTS

We wish to thank Drs. Dyanne Brewer and Armen Charchoglyan for their assistance with mass spectrometry analysis of cannflavins, as well as the NIMH Psychoactive Drug Screening Program (NIMH-PDSP) for help collecting the Tango GPCR assay results presented in this study. This work was supported by a Natural Sciences and Engineering Research Council of Canada (NSERC) Discovery Grant (401389), the Canadian Foundation for Innovation (CFI, 037755), and generous start-up funding from the University of Guelph (all to J.L.). A.W.M was supported by the University of Guelph (Graduate Tuition Scholarship).

## LIST OF ABBREVIATIONS

Arc: activity-regulated cytoskeleton-associated
BDNF: brain-derived neurotrophic factor
CREB: cAMP-response element binding protein
GPCR: G protein-coupled receptors
Mapk: mitogen-activated protein kinase
mTOR: mammalian target of rapamycin
PI3K: phosphatidylinositol-3 kinase
THC: Δ^9^-tetrahydrocannabinol
TrkB: tropomyosin receptor kinase B

## SUPPLEMENTARY INFORMATION

**Supplementary Figure 1.**
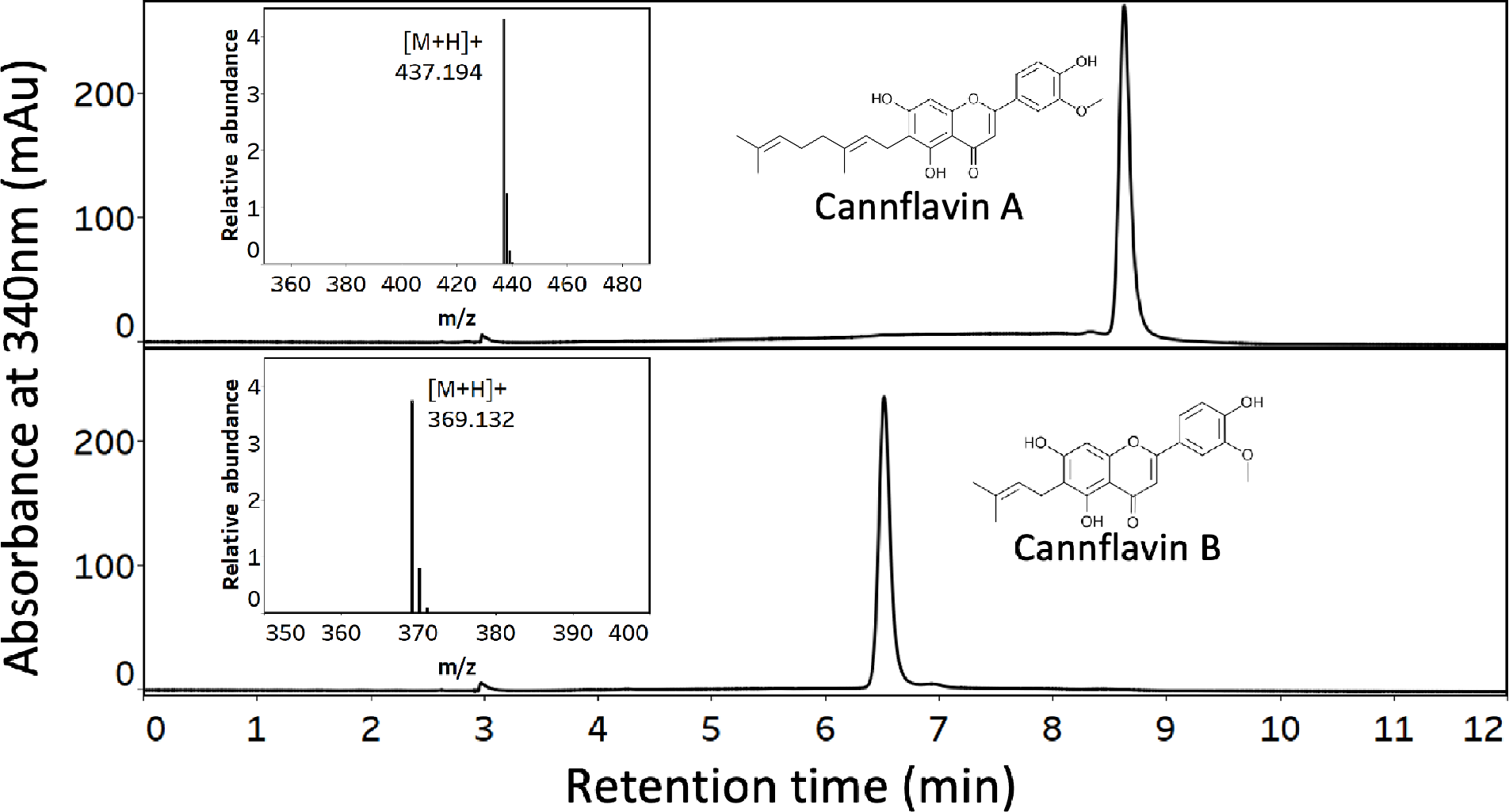
Enzymatic production of cannflavin A and cannflavin B. Representative chromatograms illustrating pooled fractions of cannflavin A (top) and cannflavin B (bottom) quantified by DAD at 340 nm. Both Q-TOF mass spectra (inset) are consistent with the expected m/z of 6-geranyl chrysoeriol and 6-dimethylallyl chrysoeriol or cannflavins A and B, respectively.

**Supplementary Figure 2.**
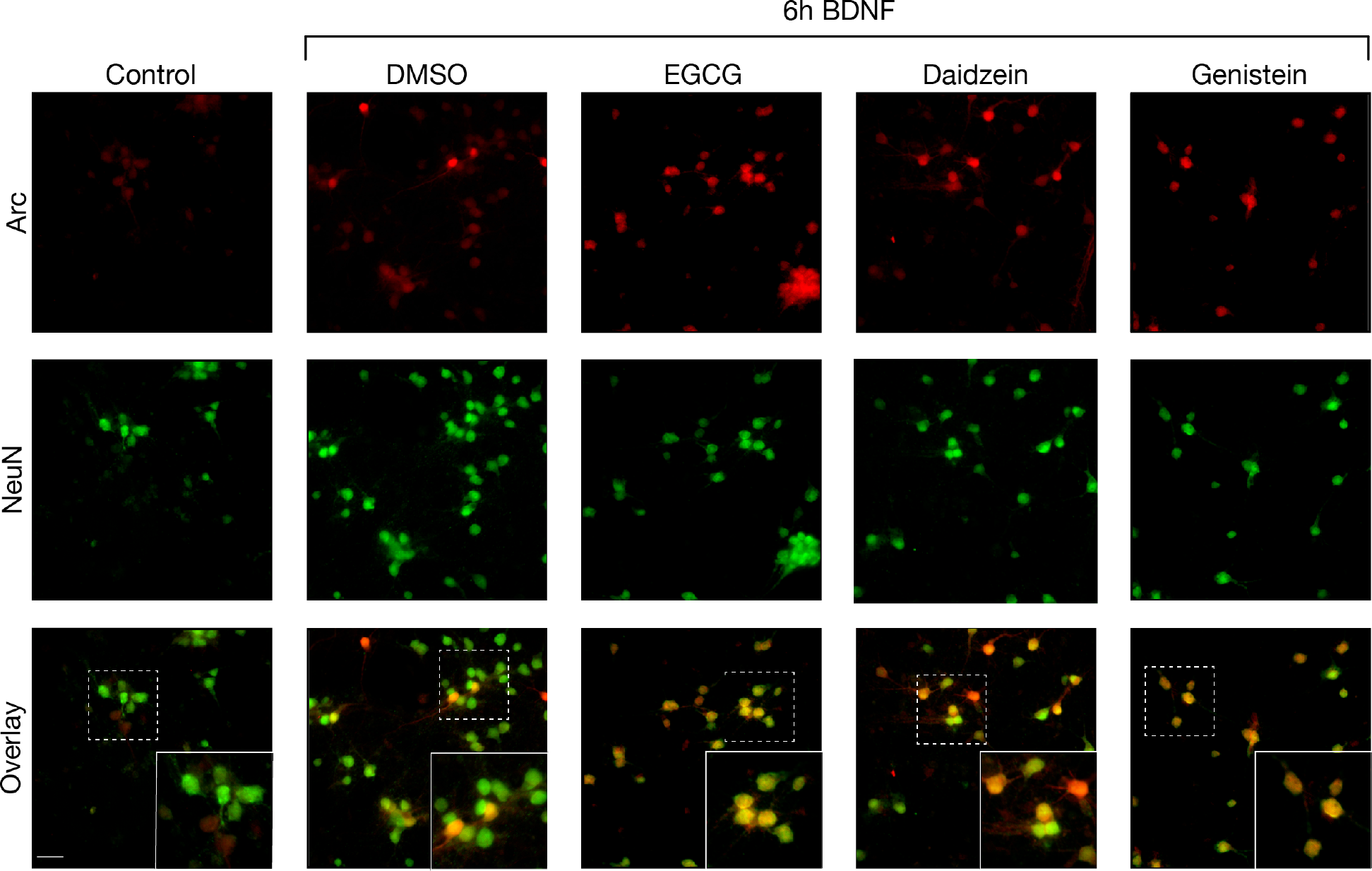
Effect of other flavonoids on BDNF-induced Arc expression level in mouse primary cortical neurons. DIV14 cortical neurons were treated with BDNF for 6 h along with ECGC, daidzein, or genistein (final concentration 10 µM). DMSO was used as control. Representative immunostaining of Arc (red fluorophore) in untreated primary cortical neurons and cells that were treated with BDNF and other compounds for 6 h. Consistent with data reported in Lalonde and colleagues (2017), daidzein and genistein increased Arc abundance at the tested concentration. Cells were co-immunostained with the neuronal marker NeuN (green fluorophore) to confirm specificity of staining to neurons. The high-magnification bottom-right insets in each panel show that Arc immunostaining in a BDNF-treated culture is particularly abundant in the nuclear compartment at the 6 h time point. Scale bar = 50 µM.

**Supplementary Figure 3.**
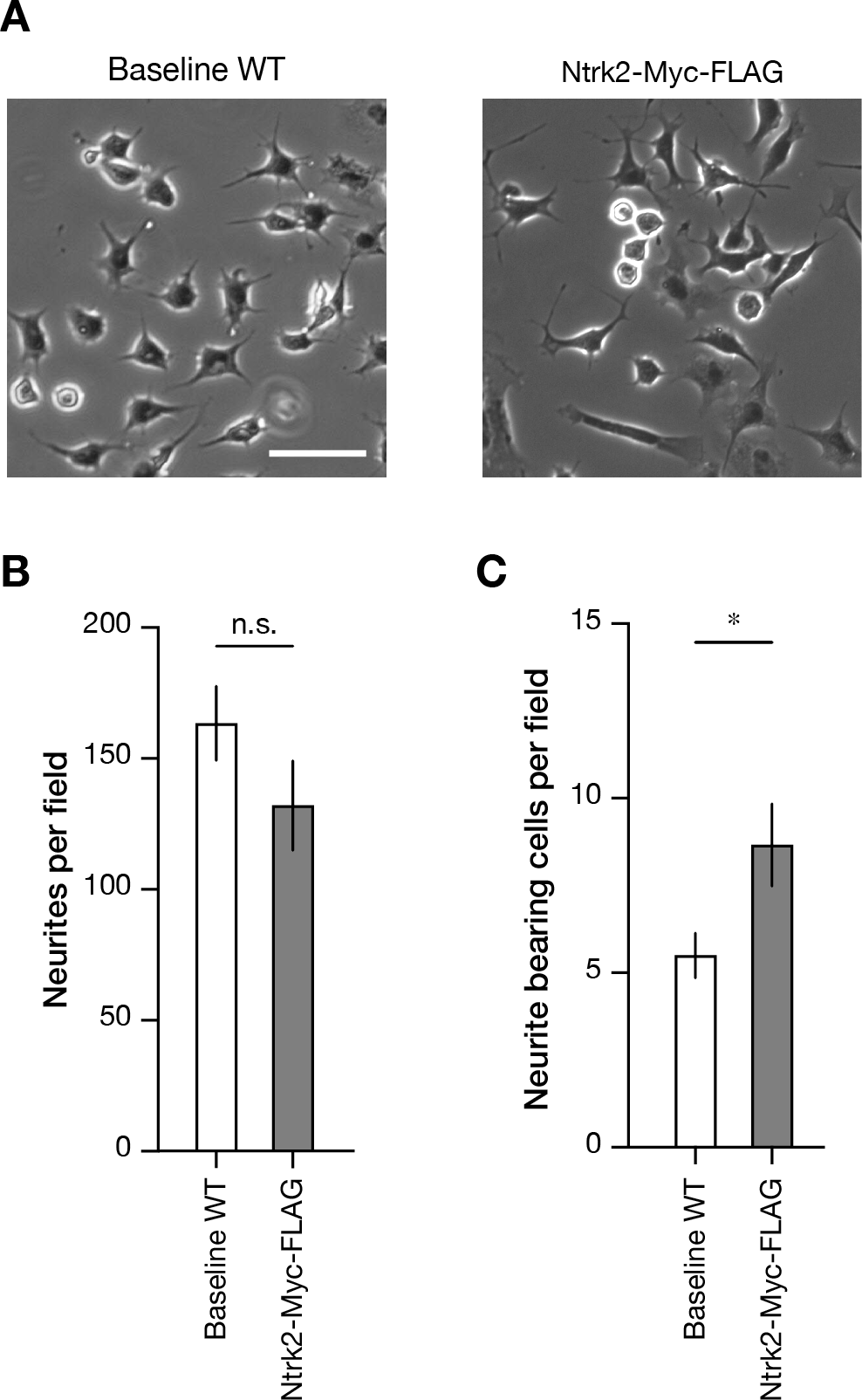
Stable overexpression of Myc-FLAG tagged TrkB increases neurite outgrowths and survival in Neuro2a cells. **A)** Phase contrast images of wild-type (WT, left) and stably overexpressing Ntrk2-Myc-FLAG (right) Neuro2a cells. Images were quantified by counting **B)** total number of neurites, **C)** total number of cells bearing neurites twice the length of cell body, and **D)** number of viable cells per imaged area. * *p* < 0.05, two-tailed unpaired *t*-test. Graphs represent mean ±SEM.

**Supplementary Table. GPCRome cannflavins report.** List of all 320 GPCRs tested in the PRESTO-Tango assay.

